# Biosynthesis cost and catabolic pathway length explain frequency of amino acid utilization as carbon sources by bacteria

**DOI:** 10.1101/2025.10.25.684513

**Authors:** Sarah F. Flickinger, Matti Gralka

## Abstract

Heterotrophic bacteria exhibit remarkable metabolic diversity. Experimental phenotyping of individual strains is often impractical due to the diversity of metabolic strategies, highlighting the need for predictive approaches based on biochemical principles. Here, we demonstrate that bacteria preferentially catabolize amino acids that are energetically inexpensive to synthesize and require shorter degradation pathways. Our results show how general biochemical constraints shape amino acid catabolism across bacterial lineages and provide a framework for integrating microbial physiology into predictive ecological theory.

## Intro

The assembly of bacterial communities is driven by their metabolic niches, which shape both their ability to grow and their interactions with each other^1^. A classic example of metabolic niches is the ability of bacteria to use different substrates as carbon and energy sources^2–5^. However, the diversity of microbial communities makes understanding the niches of each individual community member unfeasible. This is because even closely related organisms can have very different metabolic niches that are additionally difficult to predict from genomic information^6–10^: closely related strains differ in their consumption profiles, and closely related compounds (e.g., various amino acids) differ in their degree of utilization, i.e., the number of strains that utilize them as a carbon source^2,11,12^.

One strategy to address this problem is to attempt to understand metabolic niches using general principles, such as patterns in substrate utilization across diverse organisms that are shaped by selective pressures on microbial metabolism rooted in biochemical constraints^11,13,14^. This strategy has the potential to connect existing insights from microbial physiology to the metabolic traits of diverse organisms on an ecological scale and provide realistic parameter distributions for state-of-the-art models of microbial community assembly^15^.

Here, we hypothesized that the degree of utilization of different carbon sources is rooted in physiological principles, focusing on amino acid catabolism. By analyzing the metabolic capabilities of 186 marine bacteria, we find that amino acid utilization as a carbon source can be explained based on the assumptions that bacteria tend to catabolize amino acids that a) are inexpensive to biosynthesize and b) require only short pathways to catabolize. Our results thus connect bacterial physiology to the distributions of traits across organisms and suggest which selective pressures shape amino acid catabolism across distant bacterial species.

## Results

Previously, we profiled the ability of 186 diverse marine bacterial isolates to grow on each of all 20 amino acids as the sole carbon source (Methods)^11^. Here, we reanalyzed the resulting binary growth phenotypes and found that the frequency with which an amino acid was used as a carbon source varied widely (Fig. 1), with some amino acids consumed by the majority of strains (*e*.*g*., glutamate being used by 77% of strains) and others by hardly any (*e*.*g*., methionine by 1%). This stark difference is not explained by the potential ATP gained from catabolism (Methods). Furthermore, we observed that amino acids that are rarely used as carbon sources are generally consumed by organisms that use many amino acids, *i*.*e*., the distribution of consumption profiles across strains is nested (quantified by NODF = 0.80, Methods). This nestedness implies a strongly non-random structure to amino acid consumption that is shaped by underlying biochemical or evolutionary constraints. This finding motivated us to form two hypotheses to explain the structure of amino acid utilization as carbon and energy sources.

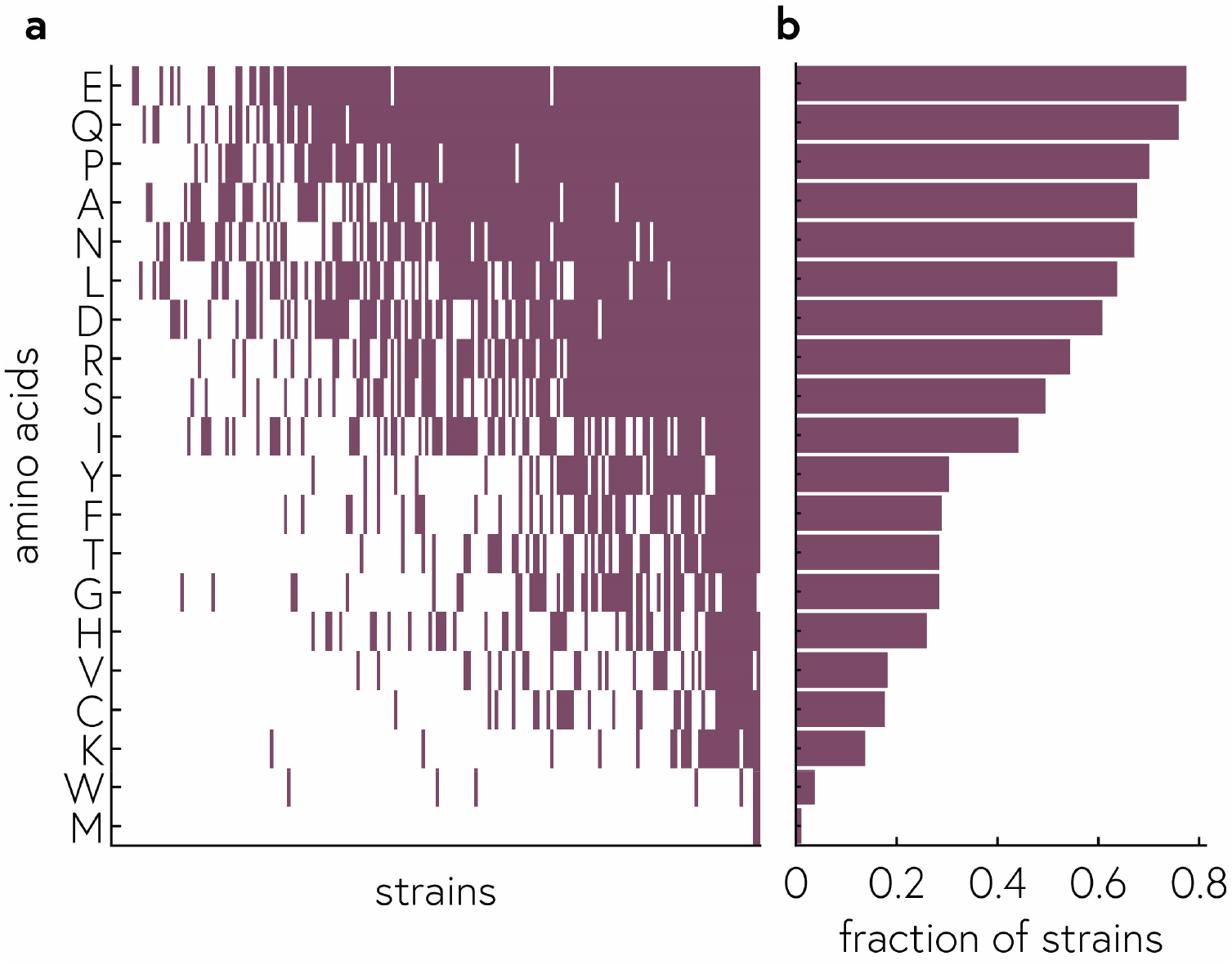

### Hypothesis 1: Bacteria tend not to catabolize amino acids that are costly to biosynthesize

Different amino acids incur different biosynthetic costs. For example, in bacteria, to synthesize glutamate from alpha-ketoglutarate takes only a single step which requires one NAD(P)H. By contrast, synthesizing methionine is a complicated process, involving multiple steps and energy carriers (ATP, NADH, etc.). The biosynthesis cost of different amino acids in bacteria was first calculated in Ref. 16. This calculation has recently been updated to include two crucial observations^17^: first, during catabolism of a carbon source some energy is liberated before the immediate precursor of a given amino acid is reached; roughly speaking, this cost should be subtracted from the biosynthesis cost. Second, the identity of the carbon source matters, since this determines how much energy is generated on the way to the precursor. We obtained the biosynthesis costs for all amino acids from Mori et al.^17^, for one glycolytic and one gluconeogenic carbon source, glucose and succinate.

Based on the differences in biosynthesis cost, we reasoned that any time a bacterium is faced with an amino acid, it is faced with the choice of using it as a building block directly or catabolizing it for energy. In our experiment, we provided only one amino acid at a time, but in their respective environments, bacteria arguably typically encounter a mix of amino acids. Therefore, we reasoned that some amino acids should be more convenient to be used for carbon and energy, not because they yield more energy, but because resynthesizing that amino acid would incur a large cost. This would lead to selective pressures to maintain the ability to catabolize energetically inexpensive amino acids but lose the ability to catabolize more expensive amino acids, which should be reserved for only those species that specialize in amino acid catabolism more generally. Therefore, we predicted a negative correlation between biosynthesis cost and the frequency of utilization across amino acids. Indeed, we found a significant negative relationship (*ρ* = −0.69, *p*= 0.0008, Fig. 2a), suggesting that catabolism of expensive amino acids is a disadvantageous and rare strategy that exists only in those species that are highly specialized in amino acid catabolism, explaining the nestedness of consumption profiles.

**Figure 2.**
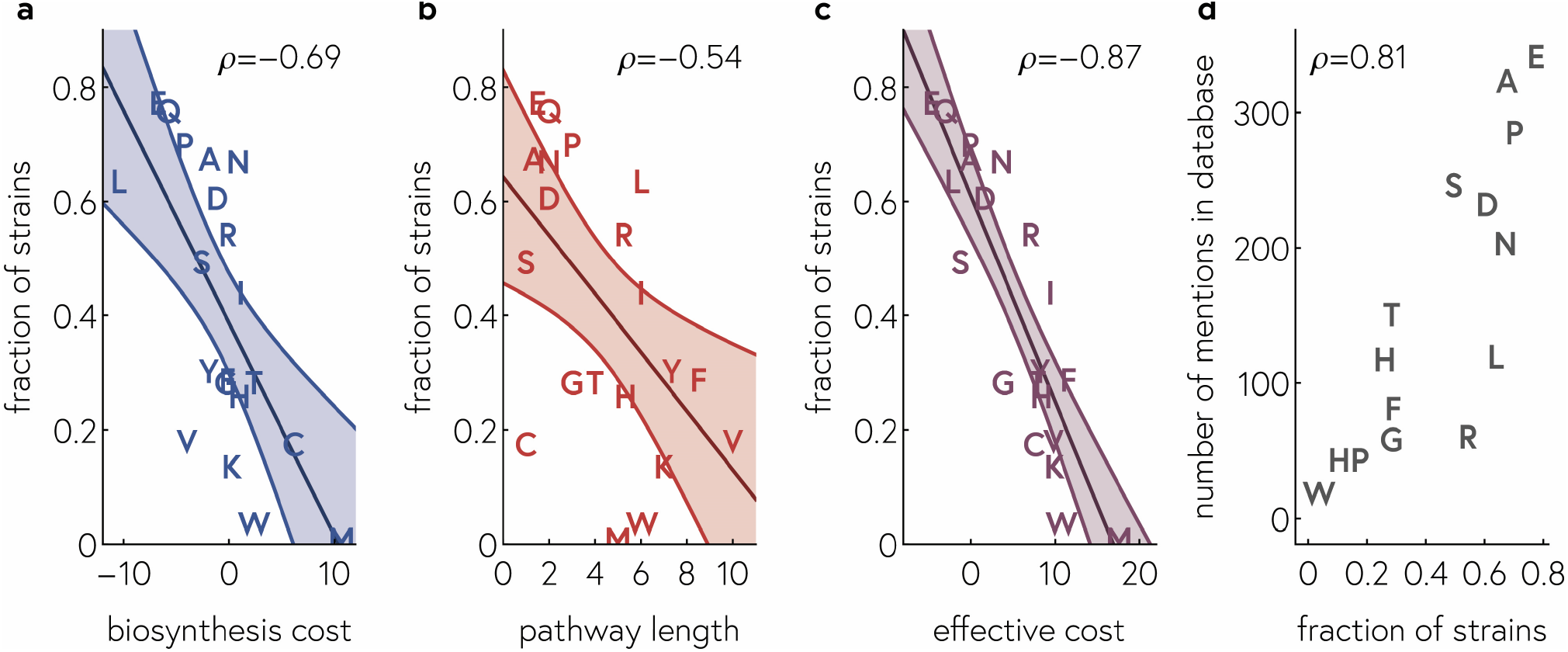
a-c) The fraction of strains growing on a given amino acid as a carbon source is correlated with the biosynthesis cost (a) and the length of its catabolic pathway (b). Taken together, the two effects produce a highly predictive model (c). d) The fraction of strains growing on a given amino acid as a carbon source in our experiments is correlated with the number of mentions of that amino acid in a large database of diverse strains.

### Hypothesis 2: Bacteria tend not to catabolize amino acids that require long catabolic pathways

Amino acid catabolic pathways vary dramatically in length, from a single step (e.g., glutamate) to 10 steps (e.g., valine). Here, we define the length of a pathways as the number of enzymes used in a catabolic pathway until the products of the pathway enter central metabolism at one of the entry points for amino acids (Methods).

We reasoned that longer catabolic pathways should incur a larger cost, both in terms of enzyme production (free proteome capacity) and, to a lesser degree, genomic maintenance (opposing genome streamlining). Thus, we predicted a negative correlation between pathway length and the frequency of utilization across amino acids which we indeed observe (*ρ* = −0.54, *p*= 0.01, Fig. 2b), indicating that longer pathways tend to be disfavored in amino acid catabolism.

### Mixed model accurately predicts amino acid utilization probability

To combine the two hypotheses, we first tested whether pathway length and biosynthesis cost were correlated and found that they were not (*ρ* = 0.05, *p*= 0.86). Therefore, we constructed a two-parameter model, accounting for biosynthesis cost and pathway length, to predict amino acid utilization across many strains. This model achieves high accuracy (*ρ* = 0.87, *p*= 4 × 10^−7^, Fig. 2c), much higher than the individual factors alone.

### Trait databases suggest the results are general

Finally, we tested whether our results might be valid more generally, i.e., across diverse strains. Previously results showed that metabolic strategies estimated from trait databases correlated well with genomic predictions^11^. Despite the difficulty in assessing the quality of the individual datasets making up these trait databases, the volume of different studies in the database means that true signals are likely to still emerge across many different labs and experiments. Therefore, we reanalyzed the database from Barberan et al.^18^ to count the occurrences of different amino acids being reported as potential carbon sources and correlated those values with the proportion of strains in our metabolic profiling experiment that utilized that amino acid. We found a strong correlation between the two datasets (Fig. 2d), suggesting that our experiments are representative of amino acid utilization patterns across thousands of diverse bacterial strains.

## Discussion

Our results show that the probability of amino acid utilization across diverse bacteria can be explained by two simple physiological principles: the biosynthetic cost of the amino acid and the length of its catabolic pathway. Together, these variables capture the main selective pressures shaping bacterial metabolic repertoires: the trade-off between conserving biosynthetically expensive building blocks and minimizing the proteomic and genomic investment required to degrade complex substrates. These findings extend classical views of metabolic niche differentiation by showing that large-scale patterns of substrate use can arise from general biochemical constraints, even in the absence of detailed genomic or regulatory information.

The strong predictive power of this physiological model suggests that microbial metabolic traits may be more structured, and thus more predictable, than often assumed. While individual species differ in regulation and ecological strategy, the overall distribution of catabolic capabilities appears to reflect fundamental constraints that transcend phylogeny. This perspective aligns with emerging frameworks in microbial ecology that treat trait distributions as outcomes of trade-offs shaped by cellular economics rather than purely by historical contingency. By identifying the biochemical basis of these trade-offs, our results provide a mechanistic foundation for predicting which substrates are likely to support growth in diverse microbial communities.

More broadly, linking physiological costs to ecological trait distributions has implications for modeling community assembly and ecosystem function. Quantitative trait models of microbial communities typically require empirical distributions of substrate use, which are often poorly constrained. Our work provides a route to derive these distributions from first principles. In this sense, the principles uncovered here offer a step toward integrating microbial physiology into predictive ecological theory - an essential goal for understanding and managing microbial processes in natural and engineered ecosystems.

## Methods

### Experiments

The main dataset is taken from Gralka et al.^11^. Briefly, 186 bacteria strains isolated from marine particles were grown on 140 individual carbon sources, including all 20 amino acids. This experiment was repeated 3 times, though not all strains were included in each replicate. Here, we scored growth on a given amino acid as a maximum OD above a threshold value of 0.08 in at least one replicate. We report the fraction of strains that grew on a given amino acid as a carbon source.

The database of traits from IJSEM was compiled by Barberan et al.^18^. For Fig. 2d, we counted the number of occurrences of each amino acid in the database.

### Biosynthesis cost C

Biosynthesis costs were taken from Mori et al.^17^. Briefly, for a given amino acid, the effective biosynthesis cost is computed as the difference between the amount of ATP gained by catabolizing the input carbon source minus the cost of biosynthesis of that amino acid. Thus, the effective cost can be negative if more ATP is liberated during catabolism (e.g., glycolysis and oxidative phosphorylation) than is required to synthesize the amino acid (e.g., glutamate from alpha-keto glutarate). Since input carbon source affects the biosynthesis cost of a given amino acid, we averaged the biosynthesis costs for glucose and succinate – one sugar and one acid, a choice motivated by our previous observation of the sugar-acid preference axis – to derive a mean biosynthesis cost across many diverse species.

### Pathway length L

Amino acid catabolic pathways were compiled from MetaCyc on Oct 13 2025. Pathways were retained if they are found in bacteria and lead to the conversion to entry point metabolites in central metabolism (pyruvate, succinate, succinyl-CoA, acetyl-CoA, oxaloacetate, or alpha-ketoglutarate). Where multiple pathways exist, we computed the geometric mean of the pathway lengths to weight shorter pathways more heavily.

### ATP gain

ATP is gained during amino acid catabolism both from the catabolic pathway itself and from the TCA cycle, depending on the amino acid, pathway, and entry point in the TCA cycle. Few complete datasets for ATP yields of amino acids seem to exist for bacteria. Here, we used the data from Wuensch et al.^19^, which provides values for 8 amino acids for a marine bacterium. For these values, the correlation of ATP yield with amino acid utilization frequency was not significant (*ρ* = 0.07, *p*= 0.86). To estimate ATP yield for all amino acids, we used values for animals^20^, which are highly correlated with those in bacteria (*ρ* = 0.86, *p*= 0.0026). Therefore, we used those values to compute the correlation of ATP yield with amino acid utilization frequency, which was also not significant (*ρ* = 0.20, *p*= 0.40).

### Effective cost

To produce Fig. 2c, we defined the effective cost as *X* = *C* + *xL*, where we computed *x* to maximize the correlation of the fraction of strains growing on amino acids with *X*. The 95% uncertainty bands in Fig. 2a-c are obtained from Mathematica’s LinearModelFit function (“MeanPredictionBands”).

**Nestedness metric based on overlap and decreasing fill (NODF)** was computed on the matrix of amino acid consumption across all strains growing on at least one amino acid, using the algorithm described in Ref. 21.

## Acknowledgements

We thank with Djordje Bajic, Frank Bruggeman, and Rachel Szabo for helpful discussions.

